# Host nutrients drive paired-substrate growth and distinct biofilm lifestyles in *Finegoldia magna*

**DOI:** 10.64898/2026.09.07.749807

**Authors:** Alison Coluccio, Francia Lopez Palomera, Susannah Lawhorn, Melanie A. Spero

## Abstract

Host-associated bacteria navigate complex nutrient landscapes where metabolites act as both growth substrates and cues that shape behavior. Yet, for most commensal and pathogenic bacteria, the nutrients and metabolisms that support persistence in the host remain unknown. *Finegoldia magna* is an obligate anaerobe that normally colonizes human skin and mucosal surfaces but also causes persistent biofilm-associated infections on implanted medical devices and in chronic wounds. Here, we developed a defined medium to investigate how nutrients influence *F. magna* physiology. We found that *F. magna* has a remarkably restricted metabolism that is specialized to use a limited set of host-relevant nutrients, including glycine, fructose, nucleosides, and betaine. Carbon-source screens showed glycine was the only substrate that supported growth as a sole carbon source. Instead, growth typically required two carbon substrates: a compatible electron donor-acceptor pair, suggesting redox balance imposes major constraints on its metabolism. To determine whether these constraints extend to host environments, we cultured *F. magna* in media derived from human chronic wound tissue. Despite its chemical complexity, *F. magna* displayed a similarly restricted metabolic profile, primarily consuming peptides, nucleosides, and betaine. These nutrients also directed biofilm behavior, with different metabolites promoting surface attachment or aggregation. Our findings show that although *F. magna* lacks metabolic flexibility, this opportunistic pathogen appears specialized to exploit host-derived products of skin physiology, tissue damage, and inflammation. This work suggests that host-associated bacteria with highly specialized metabolisms may be especially responsive to nutrient availability, linking local metabolite composition to key persistence behaviors like biofilm formation.

**Importance:** Nutrient availability is an important environmental cue regulating bacterial behavior, yet these relationships remain poorly defined in host-associated bacteria. This is particularly true for obligate anaerobes, which dominate the human microbiota, but whose metabolic requirements are largely unknown and difficult to infer from genome content. Here, we show that the skin commensal and opportunistic pathogen *Finegoldia magna* has a highly restricted metabolic repertoire, typically requiring two compatible carbon substrates for growth due to redox balance constraints. By culturing *F. magna* in media derived from human chronic wound tissue, we show that this metabolic specialization extends to complex host environments, where it mainly consumes peptides, nucleosides, and betaine. These same metabolites drive biofilm behavior, altering both surface attachment and aggregation. Our findings suggest that host-associated bacteria with a high degree of metabolic specialization may be particularly sensitive to local nutrient availability, wherein host-derived nutrients shape critical persistence strategies like biofilm formation.

## Introduction

For host-associated bacteria, the nutrient landscape serves as both a source of essential resources and a set of environmental cues that shape behavior. Because nutrient availability reflects host physiology and the activities of nearby microbes, bacteria alter their behavior in response to specific metabolites (1–3). In bacterial pathogens, metabolism is often tightly coupled to virulence. The presence or utilization of specific nutrients has been shown to influence immune evasion, stress adaptation, biofilm formation, and expression of virulence factors (4–7). Yet for most host-associated bacteria, particularly those that transition between commensal and pathogenic lifestyles, we know little about how nutrient utilization supports persistence or regulates behavior.

*Finegoldia magna* is a Gram-positive obligate anaerobe that commonly colonizes human skin and mucosal surfaces (8). In addition to its role as a commensal, *F. magna* is an opportunistic pathogen that causes acute skin and soft tissue infections, including severe conditions like necrotizing fasciitis (9). *F. magna* is also a prevalent member of polymicrobial chronic infections, including chronic wound infections and bacterial vaginosis, where its role in disease is unclear (9–11). Additionally, *F. magna* readily forms aggregate biofilms in vitro (12, 13), participates in multispecies biofilms in vivo (14), and adheres to ex vivo human skin (13). *F. magna* is often found in clinical contexts where biofilm growth underlies chronicity, including chronic wounds, bone infections, and infections of implanted medical devices such as stents, prosthetic heart valves, and joint prostheses (11, 12, 15–23). Given that biofilms are highly resistant to host immune defenses and tolerant to antibiotics (24), biofilm regulation has been extensively studied in several model facultative anaerobes (25, 26). However, little is known about biofilm regulation in host-associated obligate anaerobes, despite the importance of biofilm formation as a persistence strategy.

Despite its clinical relevance, the basic physiology and metabolism of *F. magna* remain poorly understood. Although several virulence factors have been previously described (27), comparatively little is known about which nutrients and metabolic activities support its growth in host environments. Prior work has shown that *F. magna* uses peptides and amino acids as primary metabolic substrates, consistent with growth via amino acid fermentation (28, 29).

Accordingly, *F. magna* has long been considered asaccharolytic, although its genome encodes the capacity for fructose catabolism (28, 30, 31). However, prior understanding of *F. magna* metabolism was derived from growth studies conducted in rich, undefined media (28), making it difficult to determine which nutrients support its growth. Defining *F. magna* metabolism will provide insight into how this organism colonizes the host, including how nutrient availability influences behaviors such as biofilm formation.

Here, we developed a defined medium to dissect the metabolic capacity of *F. magna*. We found that this anaerobe requires few exogenous vitamins and metals and relies on a very narrow range of carbon substrates. Even among these carbon substrates, *F. magna* typically required compatible substrate partners to achieve robust growth, suggesting this organism has a specialized approach to maintaining redox balance during fermentation. Importantly, this restricted metabolic profile extends beyond defined laboratory conditions. Even in a chemically complex medium prepared from human chronic wound tissue, *F. magna* consumed a similarly limited set of substrates, including peptides, nucleosides, and betaine. We further show that host-relevant nutrients drive distinct biofilm behaviors, with different metabolites favoring surface attachment or aggregation. These findings reveal how a metabolically specialized anaerobe is well-adapted to exploit host-derived nutrients, providing a framework for understanding how nutrient requirements shape bacterial growth and behavior in host environments.

## Results

### Defined medium identifies two-substrate fermentation in *F. magna*

To better understand the metabolic capacity of *F. magna* and how nutrient availability influences its physiology, we developed a defined medium (DM) for this organism. As a starting point, we modified a DM previously developed for the anaerobe, *Clostridium sporogenes* (32). The DM consists of basal salts, amino acids, vitamins, fatty acids (Tween), and trace metals (Tables S1 – S5). Initially, we cultured *F. magna* in an anaerobic glove box with an N_2_/H_2_ atmosphere. Under these conditions, *F. magna* exhibited robust growth in rich medium (Todd Hewitt Broth supplemented with Tween 80 (THB-Tween)) and in DM with glycine, but not fructose, as a sole carbon source (Fig. 1A). Its growth on glycine was expected since *F. magna* was previously shown to grow via glycine fermentation (33). However, *F. magna* also encodes genes for fructose transport and metabolism (31), so its weak growth on fructose has remained a longstanding and unexplained observation.

**Figure 1:**
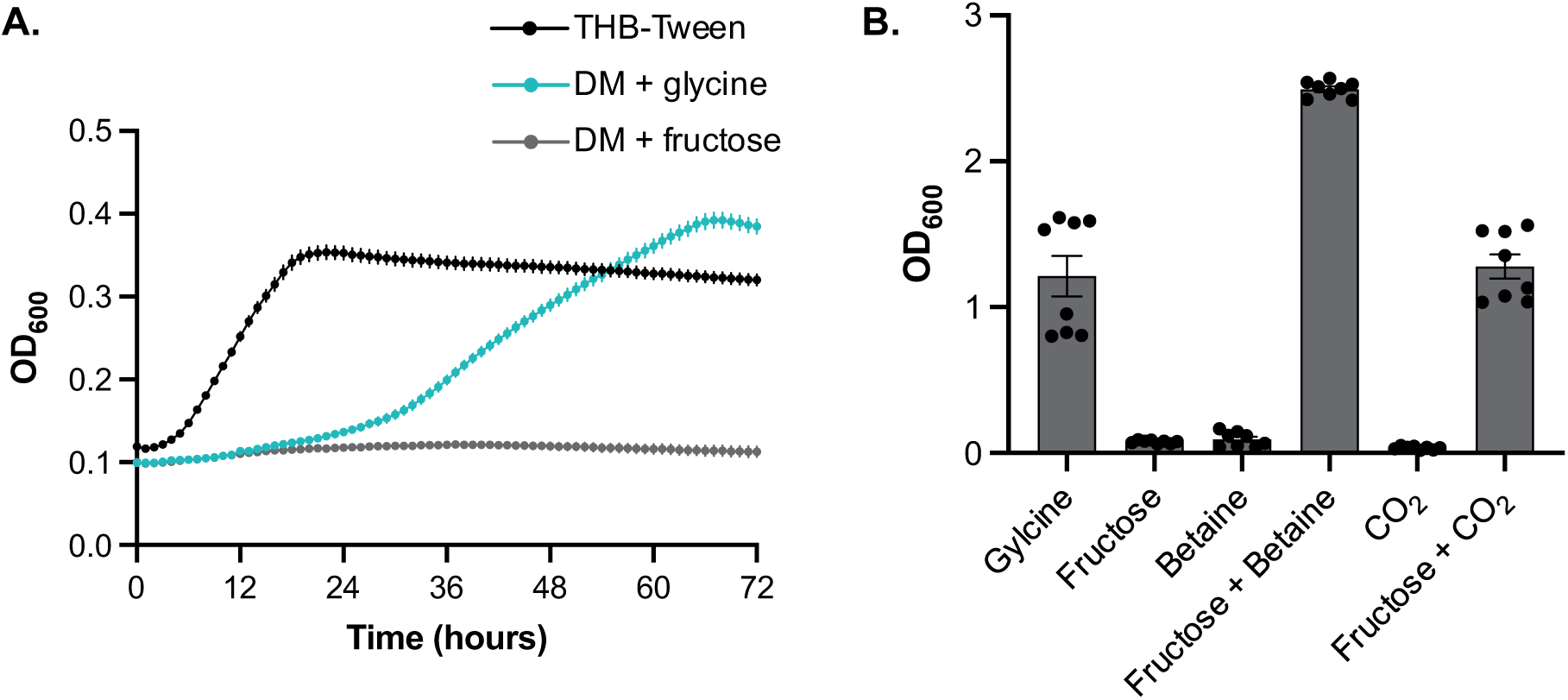
Defined medium shows *F. magna* requires two substrates for growth with fructose. **A.** Growth curve (OD_600_ over time) demonstrating that *F. magna* exhibits robust growth in rich undefined medium (THB-Tween) and in the defined medium (DM) when glycine, but not fructose, is provided as the growth substrate. Data show the mean ± standard error of the mean (SEM) of 3 replicates from a representative experiment. **B.** Final cell density (OD_600_) of *F. magna* cultures after 72 hours of growth in DM supplemented with different substrates. *F. magna* is able to grow with fructose when it is paired with betaine or CO_2_. Data show the mean ± SEM for 8 replicates across 2 independent experiments.

We hypothesized that fructose utilization might require a second substrate to maintain redox balance. Many bacteria grow via two-substrate fermentation, wherein one substrate serves as an electron donor while a second acts as an electron acceptor. This strategy is commonly used by anaerobes to conserve energy while maintaining redox balance (34–36). Consistent with this hypothesis, *F. magna* grew robustly in DM supplemented with both fructose and betaine, whereas neither compound alone supported growth (Fig. 1B). Betaine serves as an electron acceptor in numerous anaerobes (37–40) and *F. magna* encodes a putative betaine reductase for catalyzing this reaction (Table S6). Next, we tested whether CO_2_ could serve as an electron sink. While CO_2_ was absent from our initial experiments, it is known to support redox balance through various pathways in other anaerobes (41–45). Indeed, *F. magna* grew in DM with fructose when also provided CO_2_, which we predict *F. magna* reduces to formate via formate dehydrogenase; formate may be further reduced, supporting electron balance and energy conservation via the reductive glycine pathway (discussed below, Fig. 4A, Table S6) (46–48). Overall, these findings show that *F. magna* growth can require paired substrates, suggesting the need to maintain redox balance restricts which metabolites support its growth.

### *F. magna* requires few micronutrients and minimal medium restricts growth of other skin bacteria

To identify which nutrients *F. magna* requires for growth, we systematically omitted amino acids, vitamins, metals, and other components from the DM. To determine whether nutrient requirements differ between metabolically distinct substrates, *F. magna* was cultured in the presence of CO_2_ with either glycine or fructose as the carbon source (Fig. 2A).

**Figure 2:**
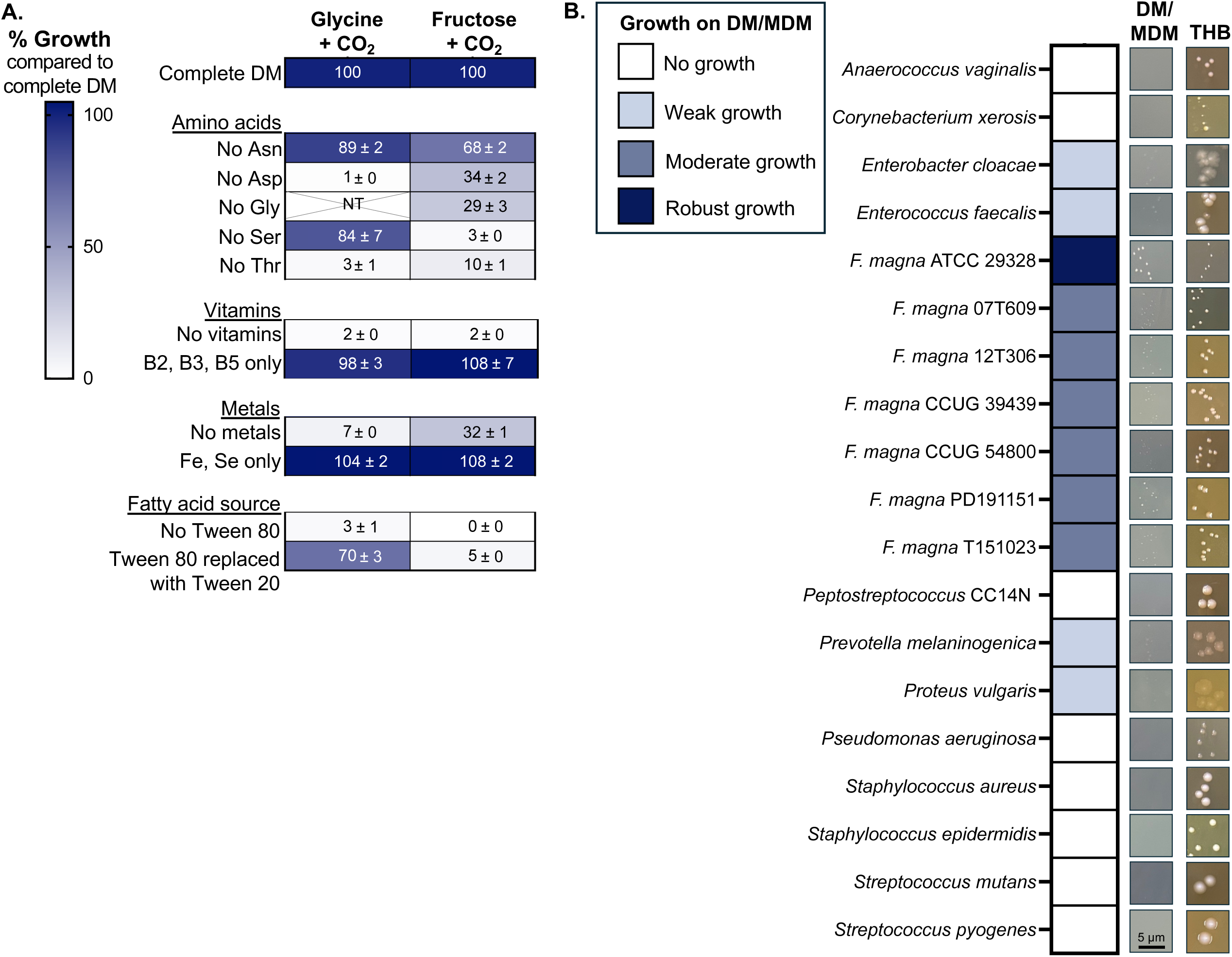
Minimal defined medium is selective for *F. magna*, which requires limited micronutrients. **A.** *F. magna* was cultured on variations of the defined medium for 72 hours and growth (final OD_600_) was recorded. Heatmap values indicate percentage growth for each condition normalized to the complete DM, which was set to 100%. Experiments were conducted with either glycine plus CO_2_ or fructose plus CO_2_ provided as growth substrates. NT = not tested. Heatmap values show the mean ± SEM from at least 8 replicates across at least 2 independent experiments. **B.** Growth on solid DM with glycine demonstrates selection for *F. magna*. Each organism was plated onto THB-Tween, DM with glycine, or minimal defined medium (MDM) with glycine. Organisms that did not form detectable colonies on DM or MDM were categorized as No growth. Organisms that formed colonies on MDM were categorized as Weak, Modest, or Robust growth when its colony diameter on MDM was <25%, 25-50%, or >75% the size of its colony diameter on THB-Tween, respectively. Representative images show each organism grown on THB-Tween (THB) and on either DM (No growth) or MDM (Weak, Modest, Robust growth).

The *F. magna* genome has predicted pathways for synthesizing five amino acids: asparagine, aspartate, glycine, serine, and threonine (31). Of those five, only asparagine was partially dispensable during growth with fructose, whereas growth with glycine did not require asparagine or serine, the latter of which can be synthesized from glycine by the reversible serine hydroxymethyltransferase reaction (Table S6) (49). Thus, contrary to genomic predictions, *F. magna* was largely unable to synthesize amino acids under these conditions. *F. magna* also required Tween for growth. Tween 80 (largely oleic acid) supported growth with both substrates whereas Tween 20 (largely lauric acid) only supported growth with glycine (Fig. 2A). This suggests that *F. magna* requires exogenous fatty acids for growth, consistent with adaptation to lipid-rich host environments like the skin (50, 51), and that the specificity of this requirement varies with carbon source availability.

In contrast, *F. magna* required a limited number of micronutrients for growth (Fig. 2A). *F. magna* failed to grow in DM lacking all vitamins, but growth was restored by the addition of only three vitamins: riboflavin (vitamin B2), niacin (vitamin B3), and pantothenate (vitamin B5), which are precursors to flavin cofactors, NAD(P), and coenzyme A, respectively. Likewise, while *F. magna* failed to grow in the absence of all metals, growth was restored by the addition of only two metals: iron (Fe) and selenium (Se). *F. magna* exhibited a more severe growth defect in DM lacking Se with glycine as the substrate, which was unsurprising given that glycine fermentation requires selenoproteins (33).

Using these results, we generated a solid minimal defined medium (MDM) containing only the amino acids, vitamins, and metals required for *F. magna* growth with glycine. We cultured various skin- and wound-relevant bacteria on DM and MDM plates to determine the selectivity of these media. Strikingly, MDM with glycine supported growth of all 7 tested *F. magna* isolates, but restricted growth of many prevalent skin and wound bacteria, including closely related anaerobes such as *Peptostreptococcus* and *Anaerococcus* (Fig. 2B). Thus, the metabolic requirements of *F. magna* differ from neighboring skin and wound bacteria, indicating that *F. magna* occupies a distinct nutrient niche within the polymicrobial communities where it is found.

### *F. magna* exhibits highly restricted carbon source utilization with limited single-substrate growth

We next sought to more broadly define the metabolic capacity of *F. magna*, including whether two-substrate fermentation represents a common strategy used by this organism. We used Biolog phenotypic microarray plates to screen for carbon sources that support *F. magna* growth either alone or in combination. We cultured *F. magna* in DM rather than MDM to maximize detection of growth-supporting substrates. To identify single substrates that can serve as a sole carbon and energy source, *F. magna* was cultured in Biolog plates using DM lacking an added carbon source. To identify substrates that support growth when paired with a compatible partner, we also screened for growth in DM supplemented with either fructose or betaine. While neither fructose or betaine support growth on their own, we propose that the combination enables growth because fructose acts as an electron donor while betaine acts as an electron acceptor (Fig. 1B). Thus, screening in the presence of fructose or betaine will identify additional metabolites that can function as electron acceptors or donors, respectively (Fig. 3A).

**Figure 3:**
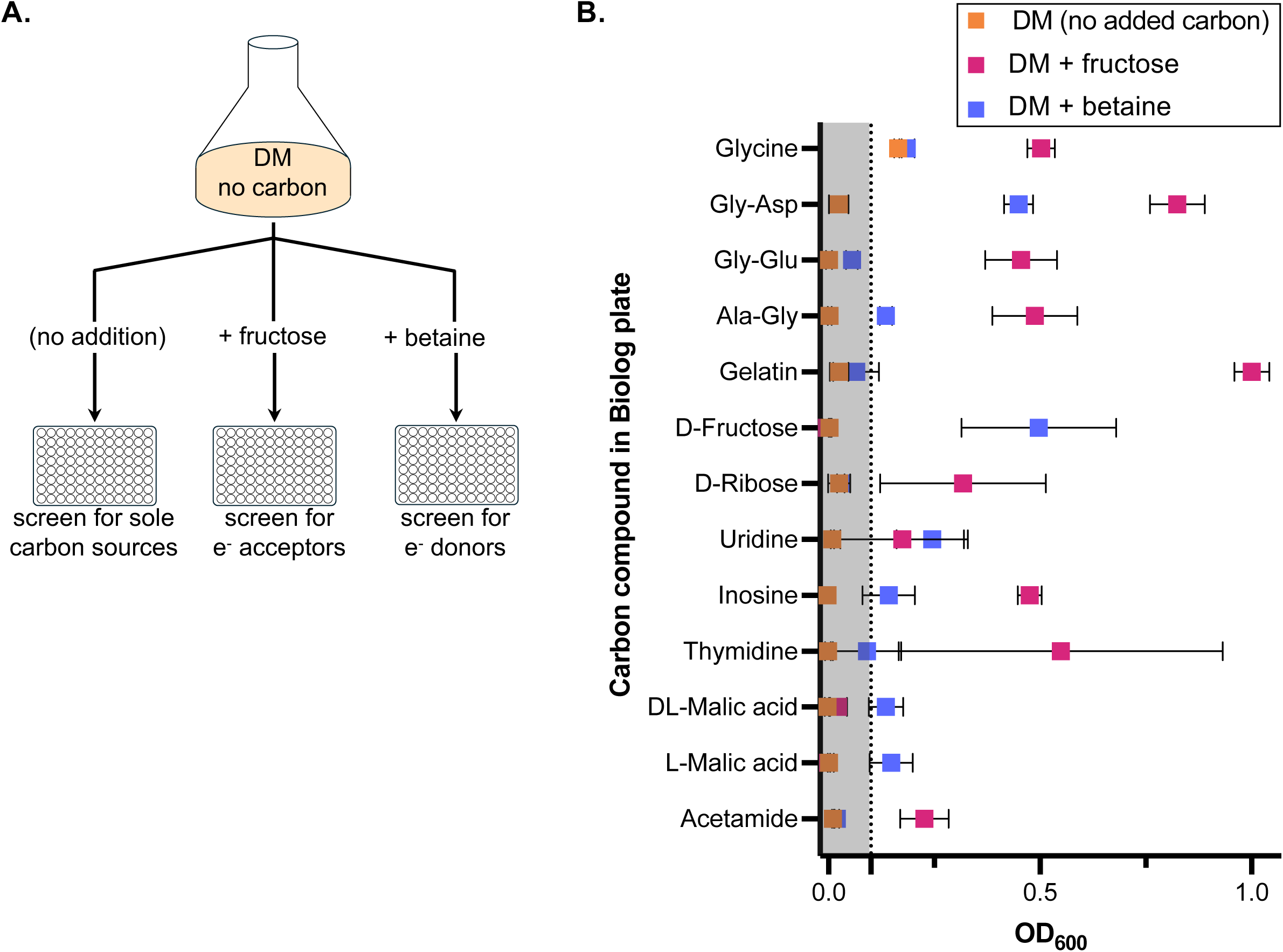
*F. magna* utilizes a narrow range of carbon substrates for growth and typically requires two substrates. **A.** Schematic of Biolog carbon source screening strategy: Biolog plates were inoculated with *F. magna* in DM lacking a carbon source, DM plus fructose, or DM plus betaine to identify substrates that support *F. magna* growth as a sole carbon source, electron acceptor, or electron donor, respectively. **B.** *F. magna* was cultured in Biolog carbon source plates for 72 hours and final cell density (OD_600_) was recorded. OD_600_ values are shown for all Biolog substrates for which at least one condition (no addition, betaine addition, fructose addition) supported growth. Growth was defined as mean OD_600_ ≥ 0.1, indicated by points to the right of the gray box and dotted line. Data show the mean ± SEM from two independent Biolog experiments.

This approach revealed that *F. magna* exhibits highly restricted carbon source utilization (Fig. 3B, Data Set S1). Of 189 compounds, glycine was the only metabolite capable of supporting growth as a sole carbon source. Instead, growth frequently required a second substrate; even under those conditions, the metabolites capable of supporting growth were limited. The addition of fructose enhanced growth with glycine and enabled growth with glycine-containing dipeptides as well as gelatin. Interestingly, fructose addition also enabled growth on ribose and ribose-containing nucleosides, such as thymidine, uridine, and inosine. The addition of betaine supported *F. magna* growth on fructose, as expected, as well as DL- or L-malic acid and several substrates identified in the fructose screen, including glycine-containing peptides, uridine, and inosine. Together, these findings show that paired substrate fermentation is a common strategy employed by *F. magna*, enabling this organism to extend its metabolic range.

### *F. magna* requires compatible electron donor-acceptor substrate pairs for growth

The observation that glycine supports *F. magna* growth as a sole carbon source is in line with glycine’s ability to act as both an electron donor and acceptor via the glycine cleavage system and glycine reductase, respectively (Table S6) (52). While this pathway is well established (33), we next used substrates identified in the Biolog screen to conduct follow-up experiments further testing our model of two-substrate fermentation as a general growth strategy for this organism. Thus, we cultured *F. magna* in DM with defined concentrations of predicted electron donor and acceptor substrates. Although pyruvate was not identified as an acceptor molecule in the Biolog screen, we investigated its role as a potential electron sink based on genomic predictions. We propose a model (Fig. 4A, Table S6) in which pyruvate formate lyase (PFL) cleaves pyruvate to form acetyl-CoA and formate. The resulting acetyl-CoA may be incorporated into biomass or converted to acetyl-P and then acetate to generate ATP. Meanwhile, we propose the resulting formate, along with CO_2_, is used to form glycine by using the glycine cleavage system in reverse (rGCS). This pathway, which is employed by other anaerobes, consumes reducing equivalents and ultimately generates acetyl-P, which likewise may be used to produce acetate and ATP (Fig. 4A) (46–48). In support of this model, growth with pyruvate required CO_2_ (Fig. S1) and we observed elevated levels of extracellular formate in cultures supplemented with pyruvate (described below, Fig. 4C).

**Figure 4:**
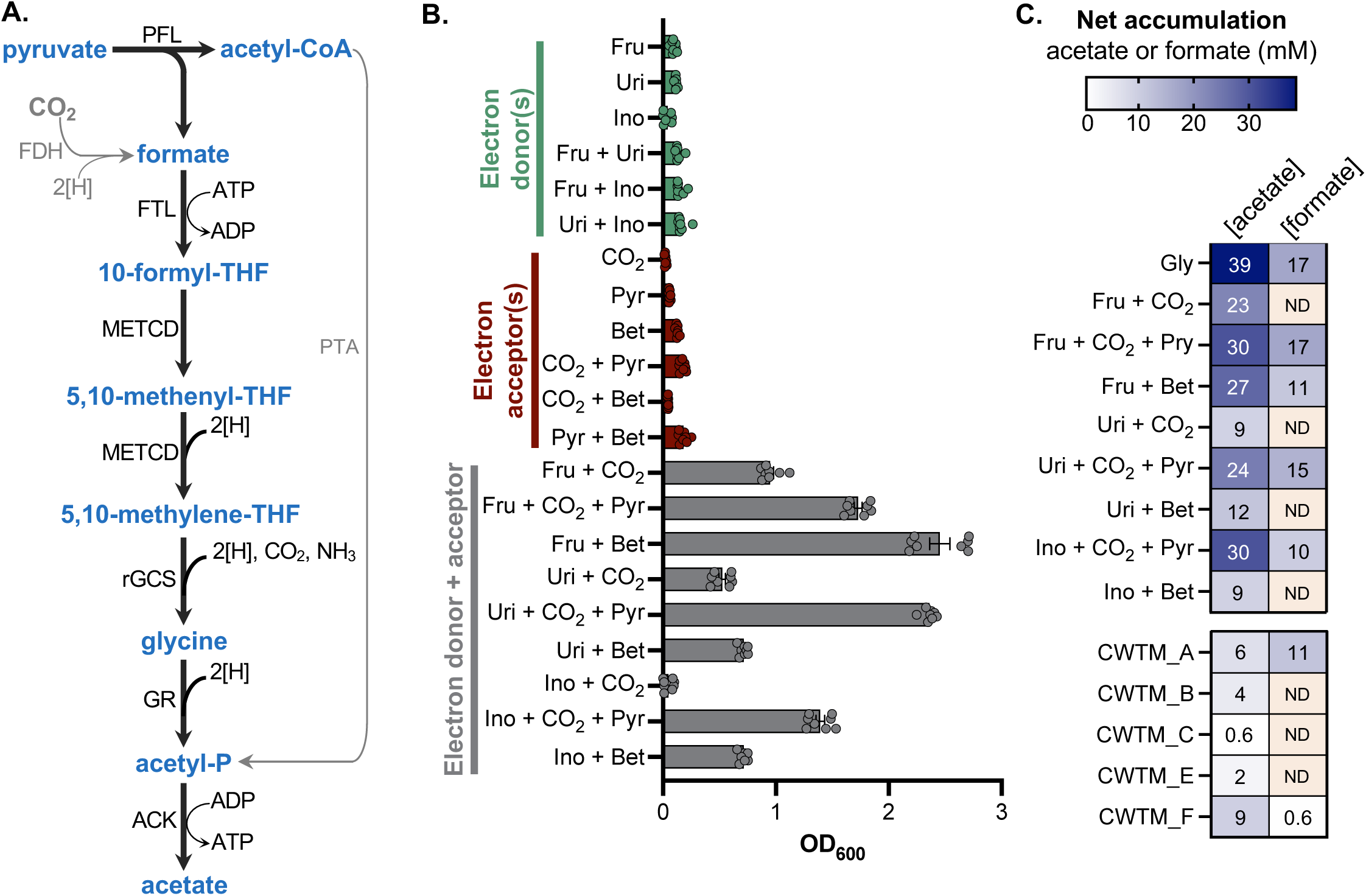
Compatible electron donor-acceptor substrates support *F. magna* growth. **A.** Proposed model of pyruvate-dependent electron consumption via reductive glycine metabolism. Pyruvate formate lyase (PFL) converts pyruvate to acetyl-CoA and formate; formate is ligated to tetrahydrofolate (THF) to form 10-formyl-THF, then reduced to 5,10-methlyene-THF, which is used by the reverse glycine cleavage system (rGCS) to generate glycine in a CO_2_-dependent reaction. Glycine reductase (GR) converts glycine to acetyl-P, which is then converted to acetate for ATP synthesis. Gray arrows indicate additional routes to formate or acetyl-P production. 2[H] denotes the consumption of reducing equivalents from an unspecified electron carrier. **B.** *F. magna* was cultured in DM with different substrates or substrate combinations for 72 hours and final cell density (OD_600_) was recorded. Data show the mean ± SEM for 8 replicates from 2 independent experiments. **C.** Heatmaps show net acetate and formate accumulation (mM) from DM or chronic wound tissue medium (CWTM) culture supernatants. DM values represent concentrations measured directly from culture supernatants because acetate and formate are absent from uninoculated DM. CWTM values were background-corrected by subtracting concentrations from the paired uninoculated control, because acetate and/or formate were detected in some uninoculated CWTM samples. ND (not detected) indicates values below the detection limit of the assay. Shown is the mean from 4 independent DM experiments or the value from one CWTM experiment. **Abbreviations:** ACK, acetate kinase; FDH, formate dehydrogenase; FTL, formate-tetrahydrofolate ligase; GR, glycine reductase; METCD, methenyl-THF cyclohydrolase/methylene-THF dehydrogenase; PFL, pyruvate formate-lyase; PTA, phosphate acetyltransferase; rGCS, reverse glycine cleavage system; THF, tetrahydrofolate; Fru, Fructose; Uri, Uridine; Ino, Inosine; Pyr, Pyruvate; Bet, Betaine.

We tested the ability of candidate electron donor or acceptor substrates to support *F. magna* growth, either alone or in combination (Fig. 4B). Fructose, uridine, and inosine were tested as electron donors, while CO_2_, betaine, and pyruvate plus CO_2_ were tested as electron acceptor substrates. As expected, *F. magna* exhibited little or no growth in DM supplemented with only a putative electron donor or acceptor. Likewise, donor-donor or acceptor-acceptor pairs did not support robust growth (mean OD_600_ < 0.18, Fig. 4B). In contrast, nearly all electron donor-acceptor combinations supported *F. magna* growth. The exception was that CO_2_ did not support growth with inosine, whereas other electron acceptor substrates did (betaine, pyruvate plus CO_2_). We also saw that the use of pyruvate plus CO_2_ as an electron sink enhanced growth on all donor substrates relative to CO_2_-only conditions. Finally, because ribose was also identified in the Biolog screen and is the sugar moiety in uridine and inosine, we tested whether ribose could similarly serve as an electron donor. Ribose addition supported modest growth when paired with pyruvate plus CO_2_ but did not support growth when paired with betaine or CO_2_ alone (Fig. S2), suggesting that nucleoside utilization is not solely driven by ribose metabolism. Taken together, these results show that *F. magna* relies on a limited set of compatible substrate combinations as a central metabolic strategy for maintaining redox balance.

To better understand how *F. magna* metabolizes its substrates, we quantified fermentation products from cell-free supernatants of cultures supplemented with different substrate combinations (Fig. 4C). Consistent with previous reports in *F. magna*, the dominant fermentation product was acetate (28, 33), which was detected in all supernatants. We also detected formate under some conditions. Formate was abundant in cultures grown with glycine, consistent with formate as a downstream metabolite of the glycine cleavage pathway (52).

Formate concentrations were also elevated in cultures supplemented with pyruvate, which we propose is generated via pyruvate formate lyase activity (Fig. 4A, Table S6). Overall, these findings demonstrate that the limited substrate repertoire of *F. magna* results in a correspondingly restricted set of fermentation products.

### Consumption of host-derived metabolites further highlights metabolic specialization in *F. magna*

To determine whether the metabolic constraints identified above also shape *F. magna* growth in host environments, we cultured this organism on media derived from human wound tissue. Chronic wound tissue medium (CWTM) was generated by homogenizing and filter-sterilizing debrided tissue from patients with diabetic foot ulcers (DFUs) or arterial insufficiency ulcers; patients were not receiving antibiotic treatment at the time of tissue collection. Five independent CWTM preparations were generated from different patient-derived samples (Fig. 5A).

**Figure 5:**
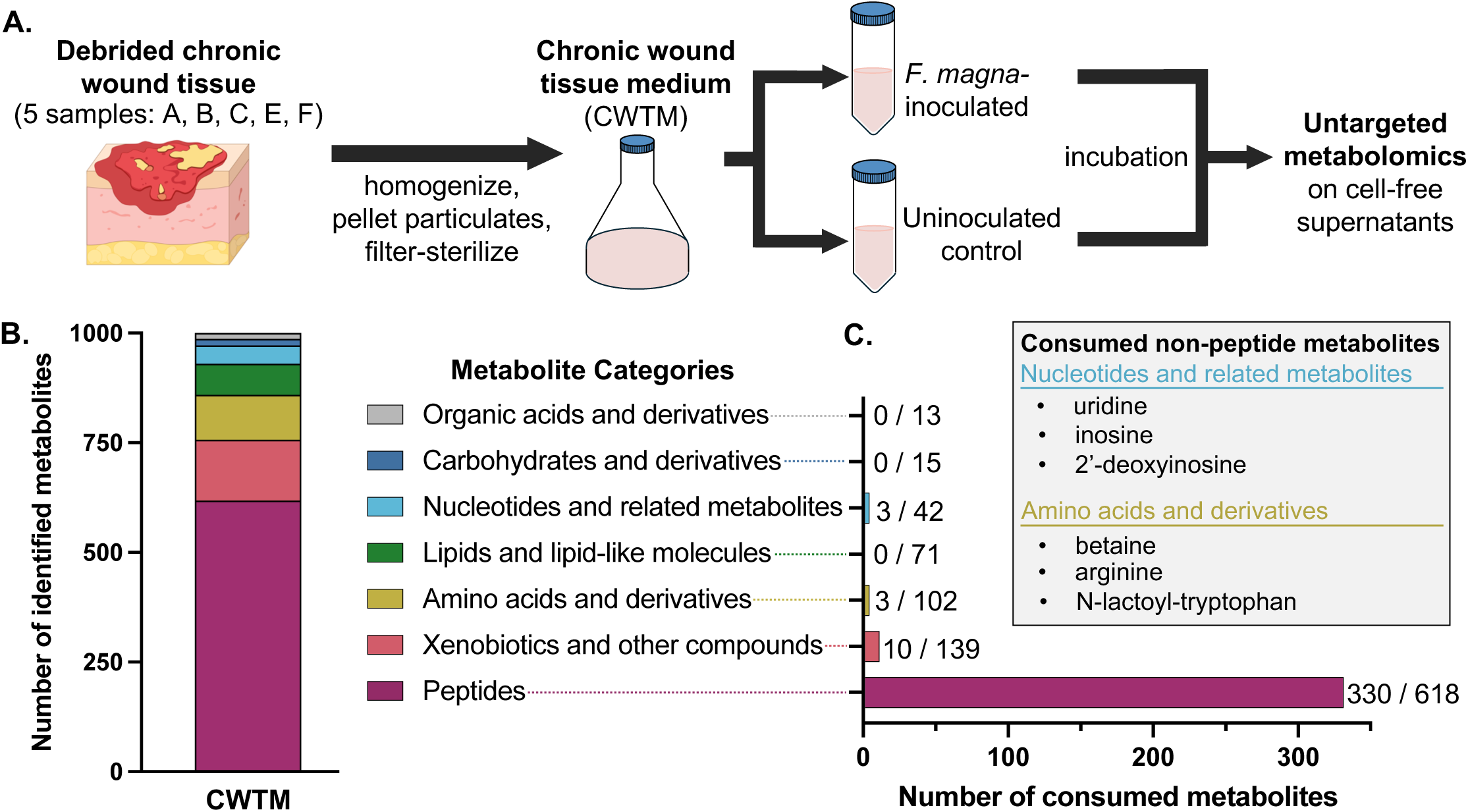
*F. magna* primarily consumes peptides from the chronic wound metabolome. **A.** Schematic of chronic wound tissue medium (CTWM) preparation and experimental design: CWTM was generated from 5 chronic wound tissue samples (each originating from 1-3 patients) and incubated with or without *F. magna*. Cell-free supernatants were analyzed by untargeted metabolomics. **B.** Metabolomics analysis identified 1,000 metabolites, and each was assigned to a metabolite category. The stacked bar graph shows the number of metabolites identified within each category from uninoculated CWTM preparations. **C.** Bars show the number of metabolites consumed by *F. magna* within each category. Values next to bars indicate the number of consumed metabolites/total number of metabolites identified in that category. The inset lists consumed non-peptide metabolites, excluding xenobiotics and other compounds. Metabolites were defined as consumed if their abundance decreased by ≥ 2-fold in *F. magna*-inoculated samples relative to matched uninoculated controls across ≥ 3 CWTM preparations.

To determine which metabolites are available in the wound environment, we performed untargeted metabolomic profiling of CWTM preparations from uninoculated control samples (Data Set S2). This approach detected 1000 metabolites representing a range of classes, including peptides, amino acids, lipids, nucleotides, carbohydrates, organic acids, and xenobiotic compounds (Fig. 5B, Data Set S3). Peptides accounted for the largest class of detected metabolites in CWTM samples (62%), consistent with elevated protease activity and extracellular matrix degradation known to occur in chronic wounds (53).

We next determined which metabolites *F. magna* consumes during growth in CWTM. *F. magna* was inoculated into each of the five CWTM preparations, alongside an uninoculated control. *F. magna* exhibited growth in all five CWTM preparations (Table S8), demonstrating these media contained sufficient nutrients for this organism. Cell-free supernatants were collected for untargeted metabolomics profiling, and metabolite abundance was compared between matched *F. magna-*inoculated and uninoculated samples for each CWTM preparation. Metabolite depletion was defined as ≥ 2-fold decrease in metabolite abundance (i.e. depleted by ≥ 50%) in *F. magna*-inoculated CWTM relative to the matched uninoculated control. To focus on metabolites that *F. magna* consistently utilized across different patient-derived samples, we defined a metabolite as consumed if it was depleted in ≥ 3 CWTM preparations. This analysis showed that *F. magna* overwhelming relies on peptides from the host metabolome, consuming over half of the peptides (330 of 618) detected in CWTM (Fig. 5C, Data Set S3).

Metabolite consumption was strikingly limited outside of peptides. Among hundreds of non-peptide metabolites detected in CWTM, only 16 were consumed by *F. magna*. These included three nucleotides (2’-deoxyinosine, inosine, uridine) and three amino acids (betaine, arginine, N-lactoyl-tryptophan) (Fig. 5C). Notably, these *F. magna*-consumed wound metabolites align with substrates identified in the Biolog screen (Fig. 3), which also found that peptides, nucleosides, and betaine serve as primary carbon substrates. Although glycine itself was not detected in the metabolomic analysis, *F. magna* consumed dozens of glycine-containing dipeptides and tripeptides. Fructose was detected in only one CWTM sample, where 30% of the metabolite was depleted. Pyruvate was detected in all five CWTM samples, but was only depleted in one sample. Finally, we quantified fermentation products from CWTM culture supernatants. Similar to findings from DM studies, only acetate and formate were detected, with acetate detected in all cultures and formate detected in two cultures (Fig. 4C). Altogether, these results show that, even within a complex metabolic landscape, *F. magna* displays a highly specialized metabolic repertoire.

### Host-relevant metabolites direct different biofilm lifestyles in *F. magna*

Because metabolites function both as resources and cues that shape behavior, we next asked whether substrate availability influences transitions between planktonic and biofilm lifestyles in this prolific biofilm-forming organism. We investigated two distinct modes of biofilm growth: surface attachment and aggregation. To quantify surface attachment, *F. magna* was cultured statically in 12-well plates, with each well containing a glass coverslip; glass-attached biofilms were quantified by staining coverslips with crystal violet after growth (Fig. 6A). Levels of surface-attached biofilm were normalized to cell density using a parallel unstained culture (OD_550_/OD_600_). When cultured with agitation, *F. magna* forms large, macroscopic aggregates (Fig. 6A). Aggregation was quantified as the fraction of cell density within the aggregate, which was calculated as 1 – (planktonic OD_600_/total OD_600_) (see methods for detail).

**Figure 6:**
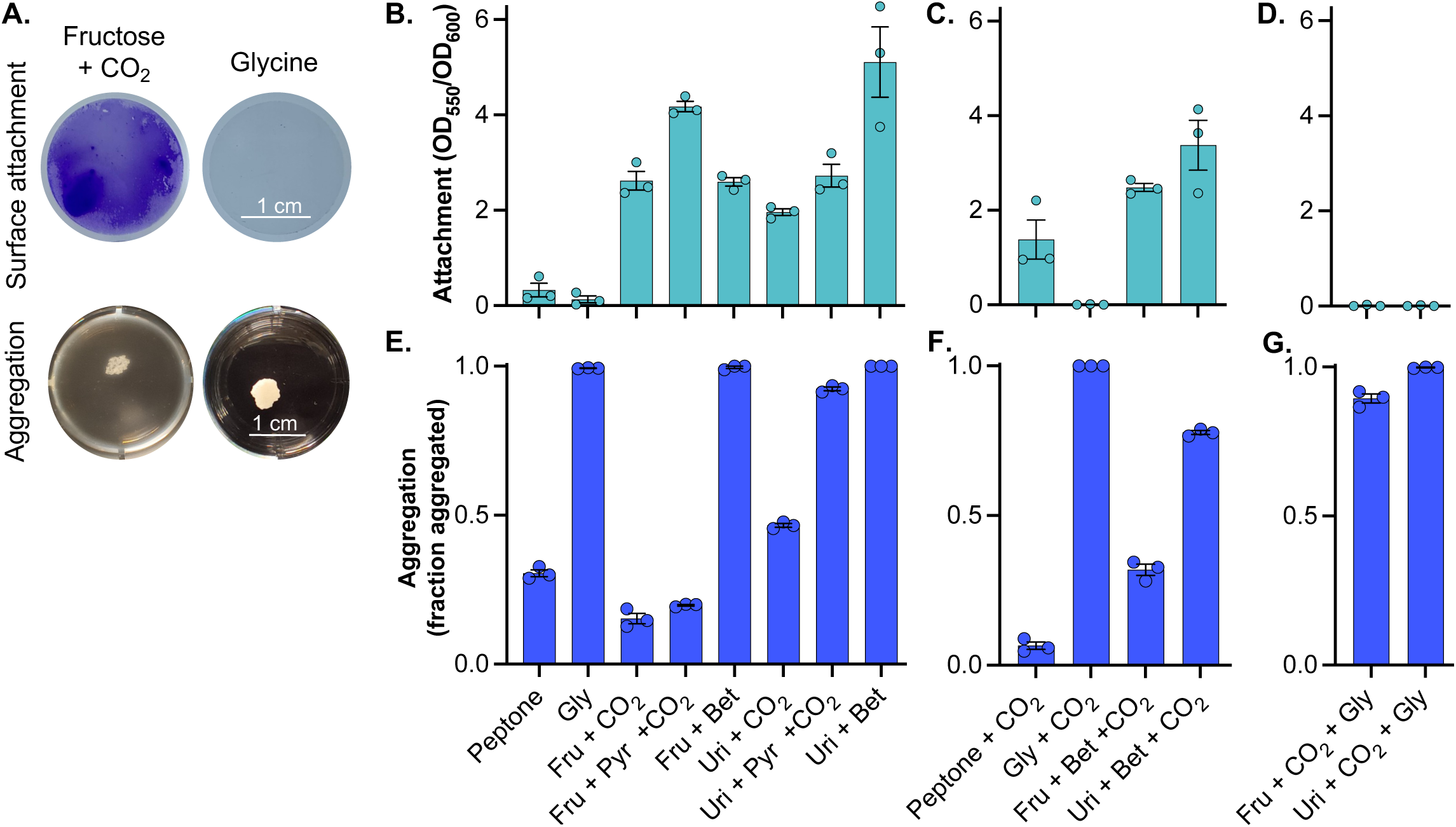
Carbon source availability drives biofilm formation in *F. magna*. **A.** Representative images of surface-attached biofilms on glass coverslips (indicated by purple crystal violet stain) and aggregate biofilms grown under two conditions: DM with glycine or DM with fructose and CO_2_. **B-D.** *F. magna* was cultured in DM with different substrates for 72 hours in the presence of a glass coverslip to promote surface-attached biofilm formation. Surface attachment was quantified as crystal violet stain per coverslip (OD_550_) normalized to total cell density (OD_600_) from parallel unstained cultures. **E-G.** *F. magna* was cultured in DM with different substrates for 72 hours with agitation to promote aggregate biofilm formation. Aggregation was defined as the fraction of cells forming aggregates, which was quantified as 1 – (planktonic OD_600_/total OD_600_). Data in panels B-G show mean ± SEM for 3 replicates from a representative experiment. **Abbreviations:** Gly, Glycine; Fru, Fructose; Uri, Uridine; Pyr, Pyruvate; Bet, Betaine.

These experiments showed that substrate availability strongly influences biofilm growth in *F. magna*, and strikingly, the carbon sources that promoted surface attachment were distinct from those that promoted aggregation (Fig. 6B-G). Interestingly, growth with peptone, which was used to mimic the peptides *F. magna* utilizes in the wound environment, resulted in relatively low levels of both surface attachment and aggregation (Fig. 6B, 6E). In contrast, glycine and fructose promoted opposing biofilm-associated phenotypes. During growth with glycine, *F. magna* did not form surface-attached biofilms but grew highly aggregated. Conversely, growth with fructose resulted in high levels of surface attachment but low levels of aggregation when paired with CO_2_ or pyruvate plus CO_2_, although *F. magna* exhibited high aggregation when fructose was paired with betaine. Similar to fructose, *F. magna* also formed surface-attached biofilms during growth with uridine, although aggregation phenotypes exhibited more variation depending on which acceptor substrate was paired with uridine (Fig. 6B, 6E).

Next, we followed up on the observation that growth with fructose resulted in low levels of aggregation in *F. magna*, except when paired with betaine. Because the low-aggregating fructose conditions included CO_2_, we tested whether the addition of CO_2_ would reduce aggregation in fructose plus betaine cultures, and indeed it did (Fig. 6C, 6F). We observed other conditions where CO_2_ addition decreased aggregation, including modest reductions in aggregation during growth with peptone plus CO_2_ or uridine plus betaine plus CO_2_ relative to their no CO_2_ counterparts. However, CO_2_ addition did not universally reduce aggregation, such as when added to glycine or uridine plus pyruvate (Fig. 6C, 6F). This suggests that CO_2_ influences aggregation indirectly, likely by altering metabolism in ways that vary depending on the broader nutrient context.

Finally, we tested biofilm phenotypes during growth in more complex mixtures, such as glycine plus fructose and CO_2_, since glycine inhibited surface attachment and promoted aggregation whereas fructose plus CO_2_ produced the opposite phenotypes. This showed that the presence of glycine was dominant, inhibiting surface attachment and promoting aggregation regardless of whether fructose plus CO_2_ or uridine plus CO_2_ were present (Fig. 6D, 6G).

Biofilm phenotypes did not correlate with cell density, which is a common driver of biofilm formation in many bacteria (54). For example, cells grown with glycine plus CO_2_ or with fructose plus pyruvate and CO_2_ achieved similar final cell densities in both assays (Fig. S3) but produced opposite biofilm phenotypes, promoting surface attachment or aggregation, respectively. Thus, nutrient conditions appear to dictate biofilm lifestyle independent of cell density, suggesting that metabolic state may drive these phenotypes. Notably, these biofilm behaviors varied in response to nutrients that *F. magna* consumes in host-derived media, including peptides, nucleosides, and betaine. These findings support a model where the host nutrient landscape directs *F. magna* towards distinct multicellular lifestyles, with distinct metabolic contexts favoring different levels of surface attachment or aggregation.

## Discussion

Host-associated microbes inhabit complex nutrient landscapes, yet the metabolic strategies enabling their persistence in host environments are poorly understood (55). While many host-associated bacteria encode flexible metabolic capacities, the skin commensal and opportunistic pathogen *F. magna* relies on few exogenous vitamins and metals and utilizes a strikingly limited metabolic repertoire. *F. magna* often required paired substrates for growth, relying on compatible electron donor-acceptor compounds to maintain redox balance. Even within complex host-derived environments, it utilized a narrow range of substrates, including peptides, betaine, and nucleosides. This degree of metabolic specialization couples nutrient availability not only to growth, but also to critical behaviors such as the transition between planktonic and biofilm lifestyles.

The identification of glycine as the only sole carbon source to support *F. magna* growth distinguishes it from other skin-resident microbes. Many skin- and wound-associated opportunistic pathogens, such as *Staphylococcus aureus*, *Pseudomonas aeruginosa*, and *Enterococcus faecalis*, utilize dozens of diverse carbon sources including carbohydrates, amino acids, organic acids, nucleotides/DNA, or fatty acids (56–62). Other skin commensals also demonstrate metabolic flexibility. *Staphylococcus epidermidis* and *Corynebacterium* spp. are capable of using glycerol, carbohydrates, amino acids, and organic acids as carbon sources (63–65). Even *Cutibacterium acnes*, a more metabolically specialized skin commensal, is known or predicted to ferment glycerol and a range of carbohydrates and related substrates (66–69). Thus, compared to other human-associated commensals and opportunistic pathogens, *F. magna* metabolism is unusually constrained.

This raises the question of why so few substrates support *F. magna* growth as sole carbon sources. Our findings suggest that redox balance requirements strongly constrain *F. magna* metabolism, such that many substrates only support growth when paired with a compatible redox partner. The use of a second substrate to achieve redox balance is a common strategy among anaerobes (34–36). However, substrate pairing requirements are largely unknown for human-associated anaerobes, outside of a few well-characterized systems.

Notably, Stickland amino acid fermentation is used by host-associated Clostridia and others (35, 36, 70, 71), and the gut bacterium *Limosilactobacillus reuteri* uses glycerol as an electron sink to enhance its growth during carbohydrate fermentation (72, 73). However, *L. reuteri* also uses numerous carbohydrates as sole carbon sources (74, 75), and many Stickland fermenters use multiple amino acid donor-acceptor pairs, with some also able to ferment carbohydrates (35, 36, 76). Thus, while two-substrate fermentation is not unique to *F. magna*, the extent to which it dominates its growth strategy is unusual. Our findings highlight that *F. magna* growth depends on the co-occurrence of compatible substrate pairs in its environment.

In light of its narrow repertoire, *F. magna* metabolism appears specialized for the skin and wound environments. Our metabolomic studies identified peptides as primary growth substrates for *F. magna* in wound tissue. Peptide-centered growth is consistent with prior reports that *F. magna* encodes numerous oligopeptide transporters and peptidases (31, 77) and is capable of degrading host proteins (78–80). In particular, *F. magna* can degrade collagen, the most abundant protein in skin (81, 82). Because collagen is glycine-rich, our finding that glycine is the only sole carbon source that supports *F. magna* growth suggests that its metabolism is well-suited to exploit skin breakdown products. Peptide utilization may be particularly advantageous in chronic wounds where elevated host protease activity promotes tissue breakdown, which might increase the availability of peptides to further benefit *F. magna* growth (53, 83). In line with the idea that host damage may create a favorable nutrient environment for *F. magna*, the abundance of this organism is enriched in chronic wounds compared to healthy skin (84, 85).

Beyond peptides, *F. magna* utilizes other skin and wound metabolites, including lipids, fructose, betaine, and nucleosides. Our DM studies found that *F. magna* requires exogenous fatty acids for growth, which has been observed in other skin commensals and is consistent with adaptation to the lipid-rich skin environment (50, 51). While lipid consumption was not detected in CWTM, this likely reflects a limitation of our methods, since untargeted metabolomics may underrepresent hydrophobic lipid species. Likewise, fructose was not readily detected by our methods, although it remains relevant to skin and wound environments because fructose-driven protein modification has been linked to extracellular matrix dysfunction, inflammation, and impaired wound healing (86, 87). Meanwhile, betaine is an osmoprotectant found in healthy skin and sweat, but it is also highly abundant in chronic wound tissue and exudate, with implicated roles in wound healing (88, 89). Finally, nucleoside utilization may also reflect adaptation to inflamed or damaged host environments. At wound sites, damaged cells and neutrophil extracellular traps (NETs) release nucleic acids into the environment (90, 91); notably, *F. magna* has been shown to activate neutrophils and induce NET release (92). Extracellular nucleotides act as inflammatory and wound-healing signals, which are further transformed by host enzymes to form nucleosides like adenosine, uridine, and inosine (90, 93). Indeed, a recent metabolomics analysis of diabetic foot ulcer tissue identified purine and pyrimidine metabolites as being associated with non-healing wounds (83). Uridine and inosine have been detected by others in chronic wound tissue, and intriguingly, these metabolites are most abundant in deeper wound tissue (88), which can be more hypoxic (94) and thus more likely to support anaerobes.

Altogether, while *F. magna* does not display broad metabolic flexibility, it appears to persist in skin and wound environments by exploiting specific host-derived substrates that are generated as products of skin physiology, tissue damage, and inflammation.

These host-relevant metabolites not only support *F. magna* growth but also direct its multicellular behavior. This finding is in line with broader work in other bacteria showing that biofilm formation and dispersal are regulated by nutrient availability, including carbon source identity, as well as metabolic or redox state (95–103). However, our findings extend this framework beyond biofilm state and abundance. While it is well established that nutrients regulate biofilm levels, far fewer studies have shown that nutrient composition directs distinct modes of biofilm formation (104, 105). In *F. magna*, different host-relevant metabolites produce distinct biofilm behaviors, promoting surface attachment, aggregation, both behaviors, or neither. The presence of peptides produced low levels of biofilm, whereas glycine specifically enhanced aggregation and uridine generally promoted high levels of both aggregation and surface attachment. Because peptides, glycine, and nucleosides reflect different aspects of skin and wound physiology, these findings raise the possibility that host nutrients cue *F. magna* lifestyle changes in different host environments. The ability to transition between these lifestyles may be physiologically important, as surface-attached and non-attached aggregate biofilms are increasingly recognized as distinct multicellular states (106–109). Both lifestyles are relevant for *F. magna*, which causes persistent infections by colonizing medical device or host surfaces (18–23) and participates in aggregate biofilms (14). For metabolically constrained organisms like *F. magna*, host nutrients may play an outsized role in driving not only growth, but also key behaviors such as organizing into multicellular communities.

Altogether, *F. magna* appears highly specialized for the host environments where it resides. By culturing *F. magna* in both defined and wound tissue-derived media, we identified an overlapping set of skin-relevant nutrients that expands this organism’s known metabolic repertoire. Thus, the defined medium captures key features of *F. magna* metabolism in a host-relevant nutrient environment, while enabling dissection of metabolic requirements. In particular, *F. magna* substrate utilization appears constrained by redox balance requirements, since robust growth typically required two substrates predicted to serve electron donor and acceptor roles.

Our findings highlight that unusual substrate combinations may be an underappreciated determinant of growth and behavior in host-associated anaerobes, whose specialized metabolic strategies may be difficult to infer from genomes or single-substrate assays (110). Such a high degree of metabolic specialization may make anaerobes particularly responsive to changes in the host nutrient environment. For *F. magna*, skin and wound products generate nutrient combinations that support growth while modulating biofilm state. Taken more broadly, these findings suggest that for metabolically specialized microbes, the host nutrient landscape may influence pathogenic potential by directing not only growth, but also persistence behaviors.

## Materials and Methods

### Bacterial strains and growth conditions

Strains used in this study are listed in Table S7. All experiments were conducted with *Finegoldia magna* strain ATCC 29328 unless otherwise specified. To culture *F. magna* anaerobically in the absence of CO_2_, cultures were incubated in a Coy Anaerobic chamber (95% N_2_, 5% H_2_ atmosphere) at 37°C. To culture *F. magna* anaerobically in the presence of CO_2_, cultures were sealed in an AnaeroPack System 2.5 L rectangular jar (Mitsubishi Gas Chemical Co, 50-25) with one GasPak EZ Anaerobe Container System Sachet (BD, 260678) and incubated at 37°C.

Overnight *F. magna* cultures were typically grown in a rich undefined medium composed of 30 g/L Todd Hewitt Broth (Research Products International, T47500) supplemented with 1% Tween 80 (Research Products International, P20390), referred to as THB-Tween. *F. magna* cells were also maintained on a rich solid medium, referred to as THA-Tween, composed of 42 g/L THB powder, 9.8 g/L sodium chloride, 0.7% v/v Tween 80, and 21 g/L bacteriological agar.

The preparation and composition of each defined medium (DM) component is described in Tables S1 – S5. To prepare 1L of DM, 250 mL of a basal salts solution was prepared fresh and combined with 25 mL of 10% sodium bicarbonate, 5 mL of 200x vitamins solution, 10 mL of 100x trace minerals solution, 5 mL of 5% cysteine, 100mL of 9% sodium chloride, 5 mL of Tween 80, 500 mL of 2x amino acids solution. Each carbon source was added from a filter sterilized 1M stock at a final concentration of 60 mM; for conditions with two carbon sources, both were present at 60 mM. The medium was brought to 1L with water, sterilized by passage through a 0.2 µm filter (Fisher, FB12566508; VWR, 76479-024) and used for experiments the same day. Carbon sources used in this study include glycine (Research Products International, G36050), D-fructose (Sigma-Aldrich, F3510), uridine (Ambeed, A135422), inosine (Ambeed, A257563), pyruvic acid sodium salt (Research Products International, P17000), betaine monohydrate (Aladdin, B106221), and D-ribose (Thermo, 132361000). Where specified, peptone (Research Products International, P20240) was added to DM at 2.6 g/L. Because inosine was not soluble as a 1M stock, inosine-containing media were prepared by adding inosine directly to the DM mixture at 60 mM, heating the medium to 60°C until dissolved, and then filter sterilizing the medium.

To prepare 1L of solid minimal defined medium (MDM), 9 g of sodium chloride, 5 mL of Tween 80, and 21 g of agar were added to 200 mL of water, which was autoclaved and then cooled to 60°C. Separately, the remaining MDM components were prepared by combining freshly made basal salts with the sodium bicarbonate, vitamins, trace minerals, cysteine, and amino acids amino acid, and carbon source (glycine) stock solutions at the same volumes used for the liquid DM; this mixture was filter sterilized, heated to 60°C, and combined with the molten agar solution. The composition of some components (vitamins, minerals, and amino acids solutions) differed between the DM and MDM, as specified in Tables S3 – S5.

### Growth experiments in DM

Overnight *F. magna* cultures were grown in THB-Tween in the absence of CO_2_. To inoculate cells in DM, overnight cultures were pelleted, washed once in phosphate buffered saline (PBS), and resuspended in PBS to an OD_600_ of 1. Cell suspensions were diluted into wells of a non-treated tissue culture plate containing prepared media to an OD_600_ of 0.005. As specified, experiments used 96-, 24-, or 12-well plates (VWR, 10861-562, 10861-558, 10861-556). For initial growth curves, 200 µl cultures with 50 µl sterile mineral oil (Macron, 6358-04) were added to 96-well plates and grown in the absence of CO_2_ in a Biotek Epoch 2 plate reader with double orbital shaking at 37°C, reading OD_600_ every 60 minutes for 72 hours. For other DM experiments, an endpoint OD_600_ measurement was recorded after 72 hours of growth at 37°C in either the presence or absence of CO_2_; endpoint experiments were performed using 2 mL cultures in 24-well plates unless otherwise specified.

Biolog carbon source utilization plates PM1 and PM2A (Biolog, 12111 and 12112) were used to screen for carbon substrates that support *F. magna* growth. For Biolog experiments, PBS-washed *F. magna* cells were resuspended in DM without an added carbon source, DM with 60 mM betaine, or DM with 60 mM fructose to an OD_600_ of 0.01 and 100 µl of cell resuspension was added to each well. Plates were incubated at 37°C in the presence of CO_2_ for 72 hours, after which an endpoint OD_600_ measurement was recorded using a Biotek Synergy H1 plate reader (Data Set S1).

### MDM selectivity assay

Preparation of solid DM and minimal defined medium (MDM), which uses 60 mM glycine as a carbon source, is described above and in Tables S2 – S5. To determine whether these media are selective for *F. magna* growth, different organisms (Table S7) were dilution plated onto DM and MDM. Each organism was initially cultured in THB-Tween for 1-2 days at 37°C under anaerobic conditions in the absence of CO_2_, except for *Pseudomonas aeruginosa* and *Corynebacterium xerosis* which were cultured in THB-Tween under aerobic conditions. Cultures were pelleted, washed once with PBS, and resuspended in PBS at an OD_600_ of 1. Cell resuspensions were dilution plated onto DM, MDM, and THA-Tween, the latter of which served as a positive control for growth. All DM and MDM plates were incubated anaerobically in the presence of CO_2_ at 37°C for 72 hours; THA-Tween plates were also incubated anaerobically in the presence of CO_2_ at 37°C for 72 hours, except for those with *P. aeruginosa* and *C. xerosis*, which were incubated aerobically at 37°C for 72 hours. After incubation, plates were imaged and the diameter of ten well-isolated colonies were quantified using the line tool function in FIJI (111). Organisms that did not form detectable colonies on DM and MDM were categorized as “No growth.” For each organism that grew on MDM, Percent Colony Size was calculated by dividing its mean colony diameter on MDM by its mean colony diameter on THB-Tween and multiplying by 100. Organisms were categorized “Weak growth” if its Percent Colony Size was <25%, “Modest growth” if its Percent Colony Size was 25 – 50%, and “Robust growth” if its Percent Colony Size was >75%.

### Chronic wound tissue medium preparation and growth assays

Deidentified chronic wound tissue specimens were obtained from Slocum Center for Orthopedics & Sports Medicine in Eugene, OR. Tissue was removed via debridement as part of routine wound care and would otherwise have been discarded. Patients were not receiving antibiotic treatment at the time of tissue collection. Tissue samples were stored at -80°C prior to medium preparation. Procedures for the collection and use of wound tissue was reviewed by the University of Oregon’s Research Compliance Services and was determined not to constitute human subjects research (STUDY00000454).

Tissue samples originated from individuals with diabetic foot ulcers or arterial insufficiency wounds. Five chronic wound tissue medium preparations were generated from tissue that was derived from one to three individuals per preparation (details provided in Table S8). To prepare the chronic wound tissue medium (CWTM), tissue was resuspended in PBS at 1 g tissue per 5 mL PBS. Tissue was homogenized via bead beating, using a BeadBug 6 Microtube Homogenizer at 8 intervals of 4,500 rpm for 30 seconds with 20 second pauses between each interval. After homogenization, samples were centrifuged at 21,300 x*g* for 60 minutes to pellet remaining particulates. The supernatant was passed through a 0.2 µm filter (Avantor, 76479-024) to sterilize the medium, which was stored at 4°C until use.

To inoculate CWTM, overnight *F. magna* cultures grown in THB-Tween were pelleted, washed in PBS, and resuspended in PBS to OD_600_ of 0.2. The cell suspension was used to inoculate CWTM at final OD_600_ of 0.01, using 120 µl cultures in 96-well plates. Cultures were incubated anaerobically at 37°C for 24-72 hours (Table S8). For each CWTM preparation, uninoculated controls were incubated in parallel.

After incubation, final cell density (OD_600_) was recorded (Table S8) and supernatants were collected for metabolomics analysis. Both *F. magna*-inoculated and uninoculated control samples were centrifuged to pellet any cells, and supernatants were passed through a 0.2 µm filter (Thermofisher, 42204-NN). Filtered supernatants were stored at -20°C or -80°C until samples were shipped on dry ice to the metabolomics facility.

### Untargeted metabolomics

Metabolite extraction and mass spectrometry analysis was performed at the Analytical & Biological Mass Spectrometry Core Facility supported by the Office of Research and Partnerships at the University of Arizona, RRID:SCR_023370.

#### Metabolite extraction

Metabolites were extracted from cell-free supernatants using methanol at an 8:1 ratio of methanol to sample volume. Following extraction, samples were split into two aliquots and dried by vacuum centrifugation. Dried aliquots were then dissolved in 70% of the original sample volume in either 20% methanol for reverse phase LC-MS or in 50% acetonitrile for HILIC LC-MS.

#### LC-MS Methods

Metabolite extracts were analyzed using liquid chromatography electrospray ionization tandem mass spectrometry (LC-ESI-MS/MS) using a Thermo Vanquish UHPLC system interfaced with a Thermo Exploris 480 Orbitrap mass spectrometer.

#### Reverse-phase C18 liquid chromatography (RP LC) tandem mass spectrometry

1 µl of metabolite extract was injected and compounds separated using a Waters ACQUITY Premier HSS T3 column (1.8 μm, 2.1 mm x 150 mm) with a column temperature of 45°C and a flow rate of 300 μL/min. Mobile phase A (water with 0.1% formic acid) and B (methanol with 0.1% formic acid) were initially 99:1, respectively. The gradient method continued as follows: 0–3 min held at 1% B; 3–19 min 1% B – 95% B; 19–20 min 95% B.

#### Hydrophilic Interaction liquid chromatography (HILIC) tandem mass spectrometry

1 µl of metabolite extract was injected and compounds separated using a ACQUITY UPLC BEH HILIC amide column (1.7 μm, 2.1 mm x 150 mm) with a column temperature of 45°C and a flow rate of 300 μL/min. The mobile phases were as follows: Mobile phase A (90% acetonitrile:10% water with 10 mM ammonium acetate with 0.1 % formic acid) and B (50% acetonitrile:50% water with 10 mM ammonium acetate with 0.1% formic acid). The gradient was identical for HILIC and RP chromatography.

#### Mass spectrometry parameters

Samples were analyzed in both positive and negative ionization modes using HCD (higher-energy collision dissociation). The HESI source parameters were set as follows: spray voltage 3.5 or 2.5 kV for positive and negative modes respectively; capillary temperature 350°C; S lens RF level 50 arbitrary units, and aux gas heater temperature 350°C. Full MS scan data were acquired at a resolving power of 120,000 FWHM at m/z 200 with the scanning range of m/z 65–975. The automatic gain control (AGC) target was set at 50%, with the maximum injection time of 100 ms. The data dependent acquisition (dd-MS2) parameters used to obtain product ion spectra were as follows: resolving power 30,000 FWHM at m/z 200, AGC target of 50% ions with maximum injection time is set to auto, isolation width 1.2 m/z, and HCD collision energies of 20%, 40%, 80%.

#### Identifications/relative quantification

For data generated, confident metabolite identifications were made using Thermo Compound Discoverer 3.4. For RP and HILIC positive and negative mode, spectra were aligned using an adaptive curve with a maximum of 0.5 min RT shift respectively and a 5 ppm mass tolerance. Peaks were selected based on a minimum intensity of 1e6 and a chromatographic S/N of 3. Detected features were grouped based on a mass tolerance of 5 ppm and a RT tolerance of 0.5. Compounds were assigned based on Isotopic pattern, RT, MS1, and/or MS2. All identifications and integrated peaks were manually validated and exported for statistical analysis.

### Metabolomics analysis

The metabolomics data set is provided in Data Set S2 and was deposited at the MetaboLights database (https://www.ebi.ac.uk/metabolights/MTBLS15488). CWTM preparations A, C, E, and F were analyzed in the same LC-MS run, whereas preparation B was analyzed in a separate LC-MS run (Data Set S2). For downstream analyses, only metabolites annotated as “Confirmed ID (High confidence)” or “Not in Internal DB. MS/MS match only” were used; low confidence annotations were excluded.

To identify metabolites that were consumed during *F. magna* growth within each CWTM preparation, peak area values for each metabolite were log_2_-transformed as log_2_(peak area + 1), where 1 was added to allow for transformation of zero values. For CWTM preparations with replicate cultures (Table S8), log_2_-transformed values were averaged across replicate *F. magna-* inoculated or replicate uninoculated controls before calculating fold changes. Next, a log_2_-fold change was calculated for each metabolite within each CWTM preparation as: log_2_(*F. magna* inoculated) - log_2_(uninoculated). Metabolites with a log_2_-fold change < -1 (i.e. >2-fold decrease in relative abundance) were defined as “depleted.” Metabolites were defined as “consumed” if they were depleted in ≥ 3 of the 5 CWTM preparations.

When multiple LC-MS features were assigned to the same metabolite, one feature was retained for final analysis. Retained features were determined via three ordered criteria: detection in the greatest number of CWTM preparations, depletion in the greatest number of CWTM preparations, and most negative log_2_-fold change across CWTM preparations. Duplicate metabolites were first removed within positive- and negative-ionization mode datasets. When positive- and negative-ionization mode data were combined, duplicate metabolites detected in both modes were again removed using the same criteria.

To characterize the composition of CWTM preparations and metabolites consumed by *F. magna* during growth, each metabolite in the final list was assigned to a chemical class: peptides, amino acids and derivatives, lipids and lipid-like molecules, nucleotides and related metabolites, carbohydrates and derivatives, organic acids and derivatives, or xenobiotics and other compounds. Chemical class assignments were manually determined using the following databases: ChEBI, HMDB, LIPID MAPS, and PubChem. The final list of nonredundant metabolites used to analyze CWTM composition and *F. magna* metabolite consumption can be found in Data Set S3.

### Ion chromatography

Fermentation products were identified from culture supernatants via ion chromatography (IC). To collect samples, cultures were pelleted and supernatants were transferred to fresh tubes, which were stored at -20°C until analysis. Samples were then thawed, diluted 1:80 (all except CWTM samples) or 1:200 (CWTM samples only) in 18 MΩ·cm water and 25 µl of sample was injected by a ThermoFisher Dionex AS-AP autosampler onto a ThermoFisher Dionex ICS-6000 ion chromatography system equipped with a Dionex Analytical EG eluent generator module, Dionex EGC 500 KOH RFIC potassium hydroxide (KOH) eluent generator cartridge (ThermoFisher, 075778), ATC-1 anion trap column (ThermoFisher, 37151), IonPac AS11-HC 4 mm analytical column and guard column set (ThermoFisher, 52960 and 52962), conductivity detector with suppressed conductivity detection using a Dionex ADRS 600 suppressor operated at 112 mA, and Dionex CRD 200 carbonate removal device (ThermoFisher, 062983). The flow rate was 1.5 mL/min. Analytes were separated using the following KOH gradient: 1 mM KOH hold for 8 minutes, followed by a linear gradient of 1-15 mM KOH over 10 minutes, a linear gradient of 15-30 mM KOH over 10 minutes and a linear gradient of 30-60 mM KOH over 10 minutes, followed by a 60 mM KOH hold for 2 minutes.

The major fermentation products detected were acetate and formate, which were identified via retention time and confirmed using standards. Standards were also run for the following metabolites, which were not detected in culture supernatants: lactate, propionate, butyrate, isobutyrate, valerate, isovalerate, hexanoate, succinate, malate, and fumarate. In addition to acetate and formate, IC methods also detected residual pyruvate in supernatants from cultures grown in DM supplemented with pyruvate, and trace pyruvate was also detected in cultures grown in DM with fructose plus CO_2_. The detection limits for acetate and formate were 56 µM and 92 µM, respectively. Because large chloride peaks were observed, DM supernatants were diluted 1:80 and CWTM supernatants were diluted 1:200 prior to IC analysis, establishing effective detection limits for the assay at 4.48 mM and 11.2 mM for acetate, and

7.36 mM and 18.4 mM for formate, respectively. Acetate and formate were absent from uninoculated DM, so net concentrations in culture supernatants were not background-corrected. However, acetate and/or formate were detected in some uninoculated CWTM controls, so net concentrations in *F. magna*-inoculated CWTM supernatants were determined after subtracting concentrations measured in uninoculated controls. To avoid overestimating acetate or formate concentrations, values below detection in uninoculated control samples were assigned the effective detection limit concentration prior to background subtraction.

### Biofilm assays

#### Aggregate biofilm assay

Overnight cultures grown in THB-Tween were pelleted, washed once in PBS, then resuspended in PBS to OD_600_ of 1.0. Cell resuspensions were diluted into 12-well plates containing prepared media at a final OD_600_ of 0.005 with a culture volume of 2 mL in each well. For growth without CO_2_, cultures were prepared and added to an anaerobic jar inside of the anaerobic glove box (95% N_2_, 5% H_2_ atmosphere), after which the anaerobic jar was removed from the glove box and placed on a shaking incubator (37°C, 120 rpm). For growth with CO_2_, cultures were prepared and added to an anaerobic jar with one sachet (described above) and placed on a shaking incubator (37°C, 120 rpm). Cultures were allowed to grow for 72 hours, then plates were removed to quantify the fraction of the culture that grew as an aggregate (fraction aggregated). First, the OD_planktonic_ of the culture was determined by transferring 1 mL of the planktonic portion of the culture (avoiding the aggregate) to a cuvette for OD_600_ measurement. Planktonic cells were then transferred from the cuvette back to the culture well, and the whole culture was vigorously pipetted to disrupt the aggregate. Next, OD_total_ was determined by transferring 1 mL of the disrupted culture to a cuvette to measure OD_600_. From this, fraction planktonic was calculated for each culture by dividing OD_planktonic_ by OD_total_. Finally, to determine what fraction of the culture was aggregated, the fraction planktonic was subtracted from 1.

#### Surface attached biofilm assay

Cell resuspensions were prepared as described above in the aggregate biofilm assay. Cells were diluted to an OD_600_ of 0.005 as 2 mL cultures in a 12-well plate, where each well contained an autoclaved circular glass coverslip (Fisher, 16004-300). Cultures were incubated at 37°C for 72 hours in the presence or absence of CO_2_. After growth, each coverslip was removed with forceps, rinsed by dipping into water, then submerged in a 0.1% crystal violet solution for 60 seconds, followed by 4 rinses in water. The coverslip was then transferred to a fresh 12-well plate and crystal violet was dissolved in 30% acetic acid for 10 minutes before measuring OD_550_ in a Shimadzu UV/Vis spectrophotometer.

## Supporting information

Data set S1

Data set S2

Data set S3

## Acknowledgements

This work was supported by grants to MAS from National Institutes of Health (R35GM155575) and a New Investigator Grant from the Medical Research Foundation of Oregon. Research reported in this publication was supported by the National Institute of General Medical Sciences of the National Institutes of Health under award number T32GM149387. We thank Wendy Johnson RN, BSN, WCC, CSWD-C and the Slocum Center for Orthopedics & Sports Medicine for their assistance obtaining and coordinating access to the deidentified debrided chronic wound tissue samples used in this study. We thank the Analytical & Biological Mass Spectrometry Core Facility at the University of Arizona for conducting the metabolomics analyses described here. We thank Lise Hald Schultz and Dr. Holger Brüggemann for generously sharing *F. magna* strains that were used in this study. We thank Sierra R. Scamfer for experimental assistance and for thoughtful discussion throughout the development of this project.

