## Supplemental Tables and Figures for "Host nutrients drive paired-substrate growth and distinct biofilm lifestyles in *Finegoldia magna*"

by

Alison Coluccio^1^, Francia Lopez Palomera^1^, Susannah Lawhorn^1^, and Melanie A. Spero^1,#^

**Table S1: Assembly of defined medium (DM)**

| **Component** | **Volume added per L DM** |
| --- | --- |
| Basal salts solution | 250 mL |
| 10% Sodium bicarbonate | 25 mL |
| 200x Vitamins solution | 5 mL |
| 100x Trace Minerals solution | 10 mL |
| 5% L-cysteine HCl solution | 5 mL |
| 9% Sodium chloride solution | 100 mL |
| Tween 80 | 5 mL |
| 2x Amino acids solution | 500 mL |
| 1 M carbon source stock solution | 60 mL |
| water | Bring to 1 L |

**Table S2: Composition of basal salts solution component used for DM preparation**

| **Component** | **Formula** | **MW (g/mol)** | **g added to 250 mL basal salts solution** | **Final concentration (mM) in DM** | **Included in MDM?** |
| --- | --- | --- | --- | --- | --- |
| Potassium phosphate monobasic | KH_2_PO_4_ | 136.1 | 2 | 14.70 | yes |
| Potassium phosphate dibasic | K_2_HPO_4_ | 174.2 | 2 | 11.48 | yes |
| Ammonium chloride | NH_4_Cl | 53.5 | 0.53 | 9.91 | yes |
| Magnesium sulfate | MgSO_4_ | 120.37 | 0.12 | 1.00 | yes |

Basal salts solution was prepared fresh and dissolved in 250 mL water during the preparation of 1 L DM; listed mM values indicate final concentrations in the complete DM.

**Table S3: Composition of 200x vitamins solution component used for DM preparation**

| **Component** | **MW (g/mol)** | **g/L in 200x vitamins** | **mM in 200x vitamins** | **Included in MDM?** |
| --- | --- | --- | --- | --- |
| Thiamine hydrochloride (B1) | 337.3 | 0.02 | 0.06 | no |
| Riboflavin (B2) | 376.4 | 0.02 | 0.05 | yes |
| Nicotinic acid (B3) | 123.1 | 0.02 | 0.16 | yes |
| D-Pantothenic Acid hemicalcium salt (B5) | 238.3 | 0.02 | 0.08 | yes |
| Pyridoxine hydrochloride (B6) | 205.6 | 0.02 | 0.10 | no |
| Cyanocobalamin (B12) | 1355.4 | 0.01 | 0.007 | no |

Concentrations shown are for the 200x vitamins solution; this solution was added at 5 mL per 1 L DM.

**Table S4: Composition of 100x trace minerals solution component used for DM preparation**

| **Component** | **Formula** | **MW (g/mol)** | **g/L in 100x trace minerals** | **mM in 100x trace minerals** | **Included in MDM?** |
| --- | --- | --- | --- | --- | --- |
| Trisodium nitrilotriacetic acid monohydrate | N(CH_2_COONa)_3_ • H_2_O | 275.1 | 1.00 | 3.64 | yes |
| Iron(II) sulfate heptahydrate | FeSO_4_ • 7H_2_O | 278.0 | 0.05 | 0.18 | yes |
| Manganese(II) chloride tetrahydrate | MnCl_2_ • 4H_2_O | 197.9 | 0.06 | 0.30 | no |
| Cobalt(II) chloride hexahydrate | CoCl_2_ • 6H_2_O | 237.9 | 0.05 | 0.21 | no |
| Calcium chloride dihydrate | CaCl_2_ • 2H_2_O | 147.0 | 0.045 | 0.31 | no |
| Zinc chloride | ZnCl_2_ | 136.3 | 0.025 | 0.18 | no |
| Copper(II) chloride dihydrate | CuCl_2_ • 2H_2_O | 170.5 | 0.01 | 0.06 | no |
| Boric acid | H_3_BO_3_ | 61.8 | 0.01 | 0.16 | no |
| Sodium molybdate (VI) dihydrate | NaMo04 • 2H2O | 241.95 | 0.045 | 0.19 | no |
| Sodium selenite | Na_2_SeO_3_ | 172.9 | 0.01 | 0.06 | yes |
| Sodium tungstate dihydrate | Na_2_WO_4_ • 2H2O | 329.9 | 0.02 | 0.06 | no |

Concentrations shown are for the 100x trace minerals solution; this solution was added at 10 mL per 1 L DM.

**Table S5: Composition of 2x amino acids solution component used for DM preparation**

| **Component** | **MW (g/mol)** | **g/L in 2x amino acids** | **mM in 2x amino acids** | **Included in MDM?** |
| --- | --- | --- | --- | --- |
| L-Alanine | 89.1 | 0.178 | 2.00 | yes |
| L-Arginine | 174.2 | 0.348 | 2.00 | yes |
| L-Aspartic acid | 133.1 | 0.266 | 2.00 | yes |
| L-Asparagine monohydrate | 150.13 | 0.3 | 2.00 | no |
| L-Glutamic acid | 183.59 | 0.368 | 2.00 | yes |
| L-Cysteine hydrochloride monohydrate | 175.63 | 0.352 | 2.00 | yes |
| Glycine | 75.1 | 0.15 | 2.00 | yes |
| L-Lysine monohydrochloride | 182.65 | 0.366 | 2.00 | yes |
| L-Histidine | 155.2 | 0.31 | 2.00 | yes |
| L-Isoleucine | 131.2 | 0.262 | 2.00 | yes |
| L-Leucine | 131.2 | 0.262 | 2.00 | yes |
| L-Phenylalanine | 165.2 | 0.33 | 2.00 | yes |
| L-Methionine | 149.2 | 0.298 | 2.00 | yes |
| L-Ornithine hydrochloride | 168.6 | 0.338 | 2.00 | yes |
| L-Proline | 115.13 | 0.23 | 2.00 | yes |
| L-Serine | 105.1 | 0.21 | 2.00 | yes |
| L-Threonine | 119.2 | 0.238 | 2.00 | yes |
| L-Tryptophan | 204.23 | 0.408 | 2.00 | yes |
| L-Tyrosine | 181.9 | 0.364 | 2.00 | yes |
| L-Valine | 117.2 | 0.234 | 2.00 | yes |

Concentrations shown are for the 2x amino acids solution; this solution was added at 500 mL per 1 L DM.

**Table S6: Predicted genes in *F. magna* metabolism**

Note: Some selenoproteins were assigned two gene IDs because the selenocysteine-encoding UGA codon can be misannotated as a stop codon.

| **Predicted gene ID** | **Locus tag (old)** | **Locus tag (new)** | **NCBI protein** | **UniProT accession** | **Predicted function** | **Predicted cofactors** |
| --- | --- | --- | --- | --- | --- | --- |
| fdhD | FMG_0290 | FMG_RS01560 | WP_012290303.1 | B0S0T0_FINM2 | formate dehydrogenase associated protein |  |
|  | FMG_0291 |  | WP_004166936.1 | B0S0T1_FINM2 | formate/nitrite transporter |  |
| grdD | FMG_0424 | FMG_RS02245 | WP_002837514.1 | B0S0U2_FINM2 | glycine/betaine reductase component |  |
| grdC | FMG_0425 | FMG_RS02250 | WP_012290397.1 | B0S0U3_FINM2 | glycine/betaine reductase selenoprotein | selenium |
| grdB | FMG_0426, FMG_0427 | FMG_RS02260 | WP_080503387.1 | B0S0U5_FINM2, B0S0U4_FINM2 | glycine/betaine reductase selenoprotein | selenium |
| grdA | FMG_0428, FMG_0429 | FMG_RS02265 | WP_080503388.1 | B0S0U6_FINM2, B0S0U7_FINM2 | glycine/betaine reductase selenoprotein | selenium |
| grdE | FMG_0430 | FMG_RS02275 | WP_004269617.1 | B0S0U8_FINM2 | glycine/betaine reductase component |  |
| trxA | FMG_0431 | FMG_RS02280 | WP_002835444.1 | B0S0U9_FINM2 | glycine/betaine reductase component |  |
| grdX | FMG_0432 | FMG_RS02285 | WP_002841250.1 | B0S0V0_FINM2 | glycine/betaine reductase component |  |
| grdA | FMG_0435, FMG_0436 | FMG_RS02300 | WP_080503389.1 | B0S0V3_FINM2, B0S0V4_FINM2 | glycine/betaine reductase selenoprotein | selenium |
|  | FMG_0437 |  | WP_002837510.1 | B0S0V5_FINM2 | thioredoxin reductase | NADP+ |
| ftl | FMG_0461 | FMG_RS02435 | WP_002838563.1 | FTHS_FINM2 | formate-tetrahydrofolate ligase |  |
| gcvT | FMG_0462 | FMG_RS02440 | WP_012290419.1 | B0S0G2_FINM2 | glycine cleavage system component |  |
| gcvH | FMG_0463 | FMG_RS02445 | WP_012290420.1 | B0S0G3_FINM2 | glycine cleavage system component | R-lipoate |
| gcvPA | FMG_0464 | FMG_RS02450 | WP_012290421.1 | B0S0G4_FINM2 | glycine cleavage system component | pyridoxal 5'-phosphate |
| gcvPB | FMG_0465 | FMG_RS02455 | WP_002838552.1 | B0S0G5_FINM2 | glycine cleavage system component |  |
|  | FMG_0466 |  | WP_012290422.1 | B0S0G6_FINM2 | glycine cleavage system component | FAD/NAD |
| metCD | FMG_0575 | FMG_RS03130 | WP_002839362.1 | B0RZW6_FINM2 | 5,10-methylenetetrahydrofolate dehydrogenase/5,10-methenyltetrahydrofolate cyclohydrolase |  |
| ackA | FMG_0852 | FMG_RS04570 | WP_002837914.1 | ACKA_FINM2 | acetate kinase | magnesium |
| glyA | FMG_0855 | FMG_RS04585 | WP_012290667.1 | GLYA_FINM2 | serine hydroxymethyltransferase | pyridoxal 5'-phosphate |
|  | FMG_1151 |  |  | B0S2H9_FINM2 | NAD-dependent formate dehydrogenase component | molybdopterin |
|  | FMG_1152 |  |  | B0S2I0_FINM2 | NAD-dependent formate dehydrogenase component | [2Fe-2S] , [4Fe-4S] |
|  | FMG_1153 |  | WP_002837310.1 | B0S2I1_FINM2 | NAD-dependent formate dehydrogenase component |  |
|  | FMG_1154 |  | WP_002837246.1 | B0S2I2_FINM2 | NAD-dependent formate dehydrogenase component | [2Fe-2S] , [4Fe-4S] |
| betT | FMG_1469 |  | WP_012291116.1 | B0S3E7_FINM2 | betaine uptake transporter |  |
| grdB | FMG_1470, FMG_1471 |  |  | B0S3E8_FINM2, B0S3E9_FINM2 | glycine/betaine reductase selenoprotein | selenium |
|  | FMG_1472 |  | WP_012291117.1 | B0S3F0_FINM2 | glycine/betaine reductase component |  |
| pflA | FMG_1494 | FMG_RS07840 | WP_012291135.1 | B0S3H2_FINM2 | pyruvate formate-lyase activating enzyme | [4Fe-4S] |
| pfl | FMG_1495 |  | WP_012291136.1 | B0S3H3_FINM2 | pyruvate formate-lyase |  |
| grdB | FMG_1606, FMG_1607 | FMG_RS08410 | WP_080503398.1 | B0S3T4_FINM2 | glycine/betaine reductase selenoprotein | selenium |
| grdE | FMG_1608 | FMG_RS08415 | WP_012291229.1 | B0S3T6_FINM2 | glycine/betaine reductase component |  |
| grdX | FMG_1609 | FMG_RS08420 | WP_041250636.1 | B0S3T7_FINM2 | glycine/betaine reductase component |  |
| pta | FMG_1615 | FMG_RS08450 | WP_012291235.1 | B0S3U3_FINM2 | phosphate acetyltransferase |  |

**Table S7: Strains used in this study**

| **Organism** | **Strain** | **Source of isolate** |
| --- | --- | --- |
| *Anaerococcus vaginalis* | ATCC 51170 | American Type Culture Collection (ATCC) |
| *Corynebacterium xerosis* | WARD470179-116 | VWR, Avantor |
| *Enterobacter cloacae* | MAS_322, isolated from human chronic wound, this study | Spero Lab, University of Oregon |
| *Enterococcus faecalis* | MAS_323, isolated from human chronic wound, this study | Spero Lab, University of Oregon |
| *Finegoldia magna* | ATCC 26328 | American Type Culture Collection (ATCC) |
| *Finegoldia magna (nericia)* | 07T609 | Brüggeman Lab, Aarhus University |
| *Finegoldia magna (nericia)* | 12T306 | Brüggeman Lab, Aarhus University |
| *Finegoldia magna* | CCUG 39439 | Culture Collection University of Gothenburg (CCUG) |
| *Finegoldia magna* | CCUG 54800 | Culture Collection University of Gothenburg (CCUG) |
| *Finegoldia magna* | PD191151 | Brüggeman Lab, Aarhus University |
| *Finegoldia magna (nericia)* | T151023 | Brüggeman Lab, Aarhus University |
| *Peptostreptococcus sp* | CC14N | BEI Resources Repository, NIAID |
| *Prevotella melaninogenica* | ATCC 25845 | VWR |
| *Proteus vulgaris* | WARD470198-316 | VWR |
| *Pseudomonas aeruginosa* | PA14 | Spero Lab, University of Oregon |
| *Staphylococcus aureus* | JE2 | Spero Lab, University of Oregon |
| *Staphylococcus epidermidis* | WARD470176-542 | VWR |
| *Streptococcus mutans* | WARD470179-170 | VWR |
| *Streptococcus pyogenes* | WARD470179-210 | VWR |

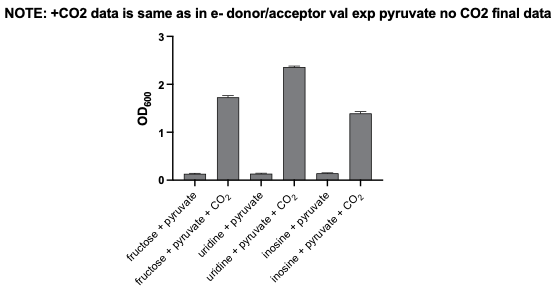

**Figure S1: Pyruvate addition does not support growth in the absence of CO_2_.** *F. magna* was cultured in DM with different substrates for 72 hours and final cell density (OD_600_) was recorded. Pyruvate addition only enhances growth in the presence of CO_2_. Data for conditions with both pyruvate and CO_2_ are the same as data shown in Figure 4; they are included here for ease of comparison. Data show the mean ± SEM for at least 6 replicates across 2 independent experiments.

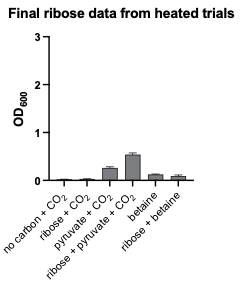

**Figure S2: Ribose supports poor or no growth as an electron donor.** *F. magna* was cultured in DM with different substrates for 72 hours and final cell density (OD_600_) was recorded. Ribose addition modestly enhanced growth when paired with pyruvate plus CO_2_, but not when paired with other substrates, suggesting it serves as a poor electron donor substrate. Data show the mean ± SEM for at least 8 replicates across 2 independent experiments.

**Table S8: Chronic wound tissue medium (CWTM) metadata**

| **Tissue ID** | **Type of wound** | **# individual patient donors** | **duration of growth incubation (hours)** | **# inoculated replicates** | **# un-inoculated replicates** | **Avg initial OD_600_, inoculated replicates** | **Avg final OD_600_, inoculated replicates** |
| --- | --- | --- | --- | --- | --- | --- | --- |
| A | Diabetic foot ulcer | 1 | 24 | 1 | 1 | 0.01 | 0.096 |
| B | Diabetic foot ulcer | 3 | 24 | 1 | 1 | 0.01 | 0.23 |
| C | Arterial insufficiency | 2 | 24 | 1 | 1 | 0.01 | 0.349 |
| E | Diabetic foot ulcer | 1 | 72 | 4 | 2 | 0.01 | 0.258 |
| F | Diabetic foot ulcer | 1 | 72 | 4 | 3 | 0.01 | 0.256 |

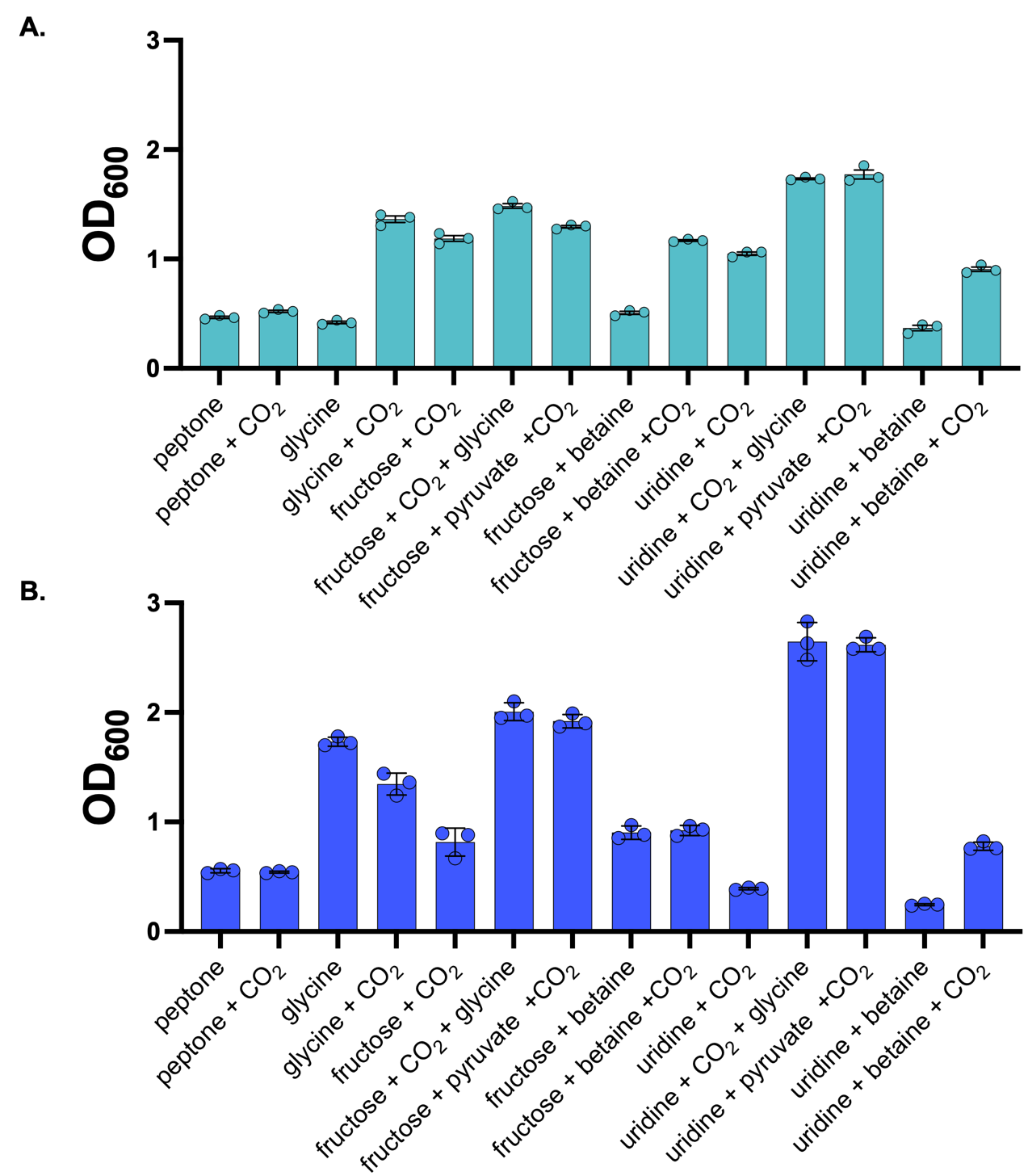

**Figure S3: Final cell densities of biofilm cultures.** Final cell densities (OD_600_) cultures analyzed in Figure 6 grown as **A.** surface attached biofilms on glass coverslips or as **B.** aggregate biofilms. These data show that surface-attached or aggregate biofilm formation do not correlate with cell density. Data show mean ± SEM for 3 replicates from a representative experiment.
